# Thermopneumatic suction integrated microfluidic blood analysis system

**DOI:** 10.1101/479386

**Authors:** Chiao-Hsun Yang, Yu-Ling Hsieh, Ping-Hsien Tsou, Bor-Ran Li

## Abstract

Blood tests provide crucial diagnostic information regarding several diseases. A key factor that affects the precision and accuracy of blood tests is the interference of red blood cells; however, the conventional methods of blood separation are often complicated and time consuming. In this study, we devised a simple but high-efficiency blood separation system on a self-strained microfluidic device that separates 99.7% of the plasma in only 6 min. Parameters, such as flow rate, design of the filter trench, and the relative positions of the filter trench and channel, were optimized through microscopic monitoring. Moreover, this air-difference-driven device uses a cost-effective and easy-to-use heater strip that creates a low-pressure environment in the microchannel within minutes. With the aforementioned advantages, this blood separation device could be another platform choice for point-of-care testing.

## Introduction

The inability to diagnose numerous diseases rapidly is a significant cause of deaths from both communicable and noncommunicable diseases in developing countries or areas with insufficient medical resources. Blood tests provide crucial diagnostic information regarding several diseases, including cancer,[1] Alzheimer disease,[2] and sepsis.[3] The gold standard process for testing a patient’s blood requires expensive laboratory equipment and well-trained technicians; however, areas with constrained resources often lack even basic diagnostic equipment and trained personnel. Furthermore, most patients are far from a clinic where laboratory services are available. Therefore, biosensors for various biomarkers, pathogens, or physiological signal detections are preferred for rapid clinical testing.[4–6]

The main limitation of typical blood tests is that relatively high volumes (in mL) of blood samples, relatively long analysis times (>1 h), and complicated processing steps are required.[7] Moreover, the reliability of testing results depends on the quality of plasma.[8] The behavior of blood cells, for example hemolysis and leukolysis, in the blood sample can affect the quality of the sample. Conventionally, blood is separated in a laboratory by using bulky and expensive centrifugation equipment that should be operated by qualified clinical technicians.[9] These drawbacks hinder the use of blood tests in areas with resource constraints.

Microfluidic technology is considered a promising approach to solve the aforementioned problems.[10–13] It miniaturizes and integrates most of the laboratory technologies into a single small chip and analyzes small amounts of samples in a short duration. Moreover, its simple operation reduces the complex multistep sample pretreatment procedures and analysis into a single step; therefore, a microfluidic system can be used by individuals without professional training. Microfluidic technology is thus crucial for achieving point-of-care testing (POCT),[14–16] which aims at performing diagnostic tests at or near a patient and at the site where care or treatment is provided.[17] Consequently, an increasing number of blood-testing devices are now based on microfluidic technology.[18–20]

Plasma separation plays a vital role in blood tests. Numerous techniques have been used to achieve plasma separation in microfluidic systems, including electroosmotic flow bifurcation (Zweifach-Fung effect),[21] geometric obstructions,[22] acoustic standing wave forces,[23] membrane filtration,[24] and cross-flow filtration.[25] However, most of these techniques have two major drawbacks, namely the complexity of the microfluidic design and requirement of an external force (provided by syringe pumps, compressed air, electropneumatic systems, high-voltage power supplies, or motors). Furthermore, treatment, such as dilution, of the blood samples before dispensing is necessary for most devices. Thus, high cost, difficulty of fabrication, low portability, and complex procedures of operation are disadvantages associated with plasma separation.[26]

A practical method for implementing on-chip flow propulsion and plasma separation was proposed by Dimov et al.[27] They demonstrated a self-contained, self-containe system that used a vacuum environment and gravitational sedimentation. The device had a relatively simple microfluidic design and working mechanism; however, the separation rate of this system was unstable. Moreover, a vacuumizer was required to create the low-pressure environment for plasma separation; this was not only time consuming and expensive but also difficult to use outside the laboratory. Therefore, we further investigated the effect of different filter trenches and microchannels on the separation rate. We varied the trench geometries, tilt angles, and the relative positions of the trench and microchannel. Consequently, we found an appropriate microfluidic three-dimensional structure that enhanced separation significantly. Moreover, to reduce the cost and improve portability, we introduced a strip heater to create a low-pressure environment for sample injection instead of a vacuumizer.

Herein, we report a gravity-driven plasma separation device with the new microfluidic design after demonstrating its separation efficiency by using a novel approach for self-containe injection (Fig. 1). Our results showed that the separation efficiency improved from 17.1% to 99.7% after adjusting the geometric design of the channel and filter trench. The production of a low-pressure environment was faster and considerably more convenient with a heater strip than with a vacuumizer. We separated the tedious blood separation procedure into two shorter steps, namely heating the air by using a heater strip for 90 s to create a vacuum-like environment and simply dispensing the blood droplets onto the inlet. The sample can be introduced and separated because of the difference in air pressure. The total separation time is <4 min, and the cost is only 1 USD for the experimental consumables. The blood separation device that we developed has two major merits, namely an extremely simple structure and operational procedure and no additional tubes or installations for external force. With these advantages, we developed a cost-effective, disposable, portable, rapid, easy-to-fabricate, and easy-to-use system, which has high potential to realize POCT in the near future.

**Fig. 1.**
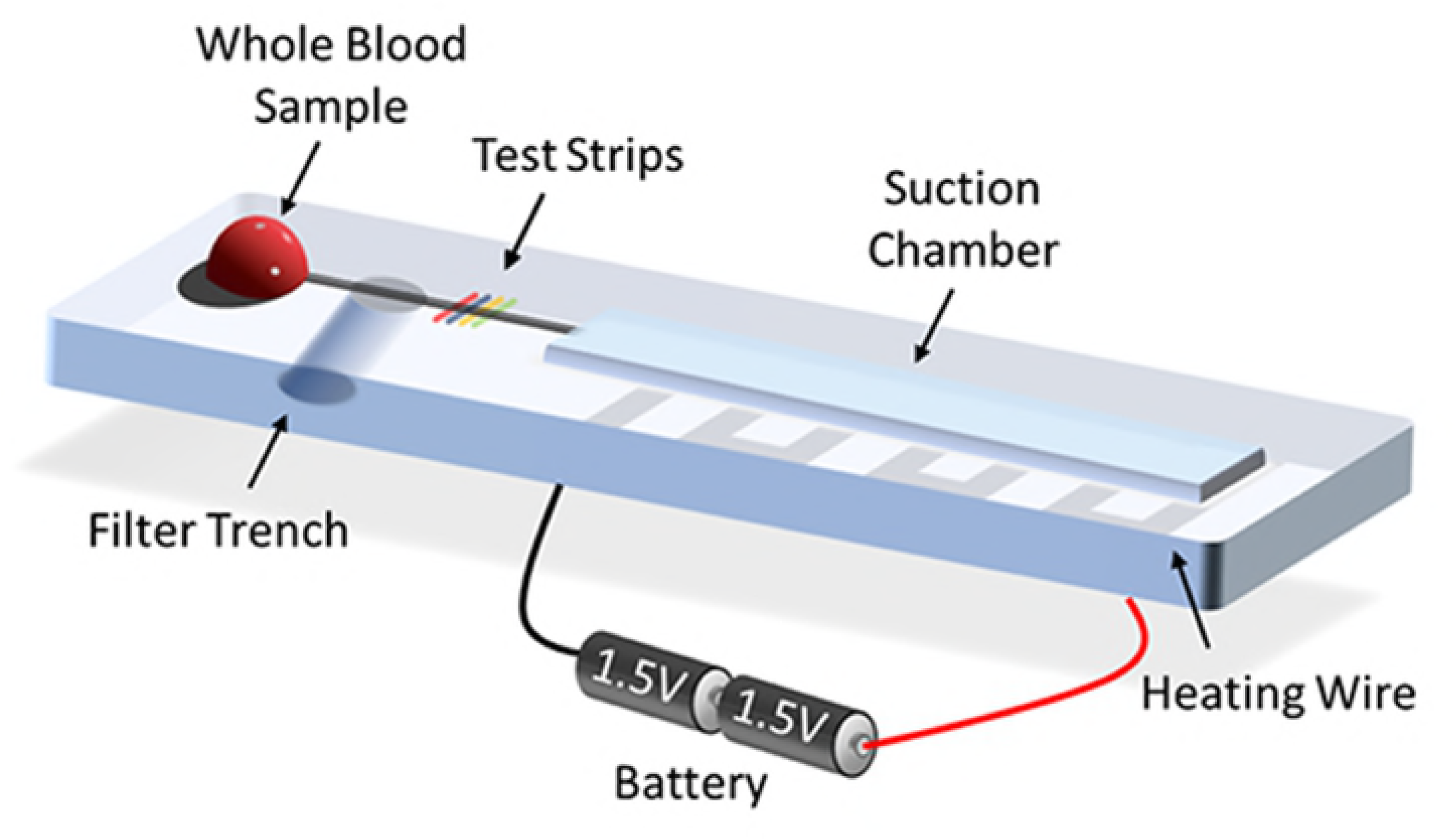
Schematic of the self-powered microfluidic device that integrates the functions of blood separation and analysis. The heating wire is set immediately under the suction chamber. Two dry cells in series constitute the power source.

## Materials and methods

### Device design and fabrication

Our microfluidic system was a PDMS-based device, composed of an inlet, a channel, a filter trench, test strips, and a suction chamber (Fig. 1). We first designed the microchannel (size: 20 mm × 1 mm × 0.1 mm) and suction chamber (50 mm × 7 mm × 1 mm) by using the Solidwork 2016 (Waltham, Massachusetts, USA) software and milled the pattern by using a computer numerical control machine (EGX-400 engraving machine, Roland, USA) to fabricate a poly(methyl methacrylate) (PMMA)-based master mold. Two 3-mm-thick PDMS (Sylgard 184 Elastomer Kit, Dow Corning Corporation, USA) slabs were made from a mixture of 8:1 (w/w). The PDMS slabs were baked at 80°C for 1 h in a precision drying oven (DOS 300, Dengyng, Taiwan). We then peeled them off the PMMA mold and punched the inlet and filter trench by using a 2-mm-diameter biopsy punch (Ted Pella Inc., USA). The PDMS slabs were irreversibly bonded to each other through infiltration of the device in oxygen plasma (Zepto Plasma, Diener, DE, USA) under 5 N oxygen pressure of 1 mbar (0.5 L h^−1^) at 60 W for 60 s. In the last step, the bottom of filter trench was sealed using Scotch Magic tape (*3M*, Maplewood, MN, USA).

### Blood samples

Human blood samples were obtained from healthy donors according to a protocol permitted by the Institutional Review Board (IRB). Fresh human blood samples were collected in a vacutainer tube from three healthy donors (BD, USA) to prevent red blood cell (RBC) aggregation. Before the experiment, we added Trypan Blue (Gibco, USA) to the whole-blood sample at the ratio of 3:100 to stain the plasma for facilitating observation. As reported in previous studies, whole-blood samples should be used within 20 min after drawing from donors. The hematocrit content of each experimental sample was adjusted to 40% through the centrifuge method.

### System operation

The power source to introduce the sample in this blood separation device is differential air pressure. In this device, we installed a heater strip immediately under the suction chamber as a method to create a low-pressure environment inside the microfluidic. Power was supplied to the heater strip by two dry cells (Panasonic, Japan) of 1.5 V in series. The experimental protocol was as follows. The air in the chamber was heated for 1 min by using the heater strip. The power was then turned off to allow the strip to cool naturally for 30 s. When the sample was loaded using a pipette, it entered the sample automatically under atmospheric pressure. With the parameters that we set, blood separation required 5 min, and detection by using urine test papers could be completed within seconds. The entire operation time of our system was <8 min, and the volume of blood samples required was only 10 μL, which can be collected directly through the finger-prick method.

### Filter trench characterization

The working principle of the separation was based on gravitational sedimentation proposed by Dimov et al.[27] We used their optimal parameter for trench diameter, which is 2 mm, and further examined how the separation efficiency would be affected by the depth, geometry of the filter trench, and the relative positions of the filter trench and channel. The device was fixed in place on a dissecting microscope (SL-730, SAGE Vision, USA) fitted with a complementary metal-oxide-semiconductor camera (700D, Canon, Japan). We designed a simple microfluidic structure for testing characterization, with an inlet, an outlet, a channel, and a filter trench. A syringe pump (NE-8000, New Era, USA) was installed at the outlet and used as the force mechanism to suck air of the channel so that the sample could be introduced. The flow rates used were 2 μL min^−1^, 1 μL min^−1^, and 0.5 μL min^−1^. Flow simulation was performed using the Solidwork Flow Simulation software.

### Microchannel design

To enhance the performance of the plasma separation, the relative positions of the filter trench and channel were further examined. The two (positions) designs tested were the buried channel and suspended channel designs. Both these channel designs were fabricated using two 3-mm-thick PDMS slabs. To fabricate the buried channel, we punched the filter trench on the channel side and bonded it with a flat PDMS slab. By contrast, the suspended channel was fabricated by punching the filter trench on the flat PDMS slab and subsequently bonding it with the PDMS slab with a straight channel.

### Image analysis and definition of separation efficiency

We recorded the results at the outlet of the filter trench in each experiment by using a charge-coupled-device camera fitted on the dissecting microscope. The recorded images were analyzed through the ImageJ software to count the pixels in the total sample and filtered plasma. A region of interest was manually identified, and the separation efficiency was calculated using the following formula:

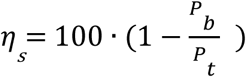

where *η*_*s*_ is blood separation ratio, *P*_*b*_ is pixel number in the area containing blood cells, and *P*_*t*_ is pixel number in the total area of plasma and blood cells that passed the filter trench. The temperature of the microfluidic device was monitored using a thermography camera (E75, FLIR System, USA). The characterization of the local temperature in space and time in the microfluidic device was collected by a built-in infracted camera.

### Analysis and detection

For the initial laboratory demonstration, we set the urine pathological changes in a patient’s blood by changing color, at the end of microchannel. Here, we demonstrated the results of variation in the glucose, protein, and pH levels as examples.

## Results and discussion

### Working principle

In this study, we demonstrated a modified filter trench system to perform blood separation by using gravity sedimentation. Previous studies have shown that gravity sedimentation has the potential to separate plasma from whole blood. Dimov et al. studied the size and proportion of blood cells in whole blood. [27] They found that blood cells are affected by the buoyancy-corrected gravitational sedimentation force (*F*_*gb*_) and fluid drag force (*F*_*d*_). Thus, the gravitational force on blood cells is significantly larger than that on plasma in a filter trench; consequently, the plasma and blood cells can be separated into an upper and a lower layer (Fig. S1). Based on this principle, we designed an extremely simple structure that required only a channel to introduce the sample to the filter trench. In our device, the filtered plasma is collected at the outlet of the trench. We developed our system based on a previous study. Furthermore, we found that the geometry of filter trench and channel design significantly affect separation efficiency.

We proposed a thermopneumatic microfluidic suction system, which is a relatively practical method for the on-chip system, as the power source of this microfluidic device. The proposed system is cost effective, simple to operate, and does not require external force mechanisms, such as syringe pumps, electropneumatic systems, or high-voltage power supplies. The suction principle is extremely simple. The fundamental principle of the thermopneumatic microfluidic suction system is based on Charles’ law, which describes the volume (*V*) of an ideal gas as being directly proportional to the absolute temperature (*T*, expressed in K) of the gas at constant pressure and number of moles:

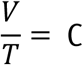

According to the ideal gas equation of state, a change in temperature affects the number of gas molecule in a fixed volume. Because PDMS has high breathability, degassing can be achieved within seconds. The cross section during the operation process is shown in Fig. 2. We designed a suction chamber at the end of the microchannel and installed a heater strip immediately under the chamber. When the heater strip is connected to the battery and the switch is turned on, the air in the chamber becomes heated within seconds, thus creating a low-pressure environment inside the microfluidic chamber. As soon as the heater strip is turned off, an inward airflow is generated by the atmospheric pressure during the process of cooling down, and the sample can be introduced automatically into the chamber. Another advantage of this mechanism is that the inward airflow can prevent the sample from being sucked back into the pipette tip, which is a common problem for low-volume sample loading.

Previous studies have also reported the use of differential pressure as the microfluidic force mechanism. However, most of them have described the use of bulky and expensive equipment to generate the vacuum or a low-pressure environment for the device. The heater strip method that we proposed is considerably less expensive and simpler than previously described equipment; however, the heater strip has high variability. The use of the heater strip reduced our cost to <1 USD.

**Fig. 2.**
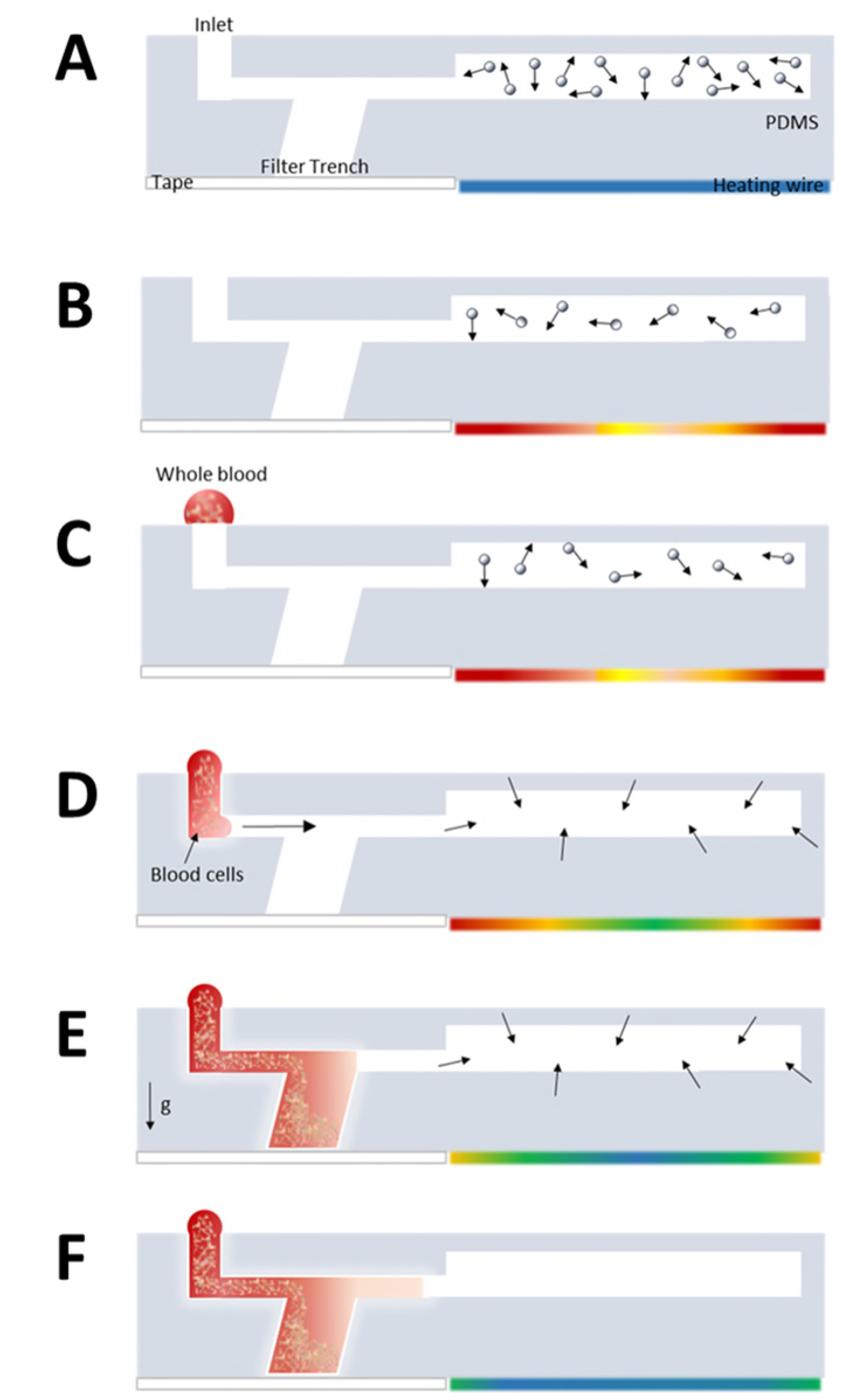
Cross section of the microfluidic device during operation: (A) original state; (B) when the heater strip starts to heat up the air in the chamber, the number of gas molecules in it decreases gradually; (C) the power is turned off and the whole-blood sample is loaded; (D) the sample is introduced into the channel through atmospheric pressure during the process of cooling down; (E) the whole blood enters the filter trench and the blood cell starts to sediment by gravity; and (F) the filtered plasma can be collected at the outlet of the filter trench. The colored strip on the bottom right side of each image (of the device) is the chromaticity bar, which indicates the temperature variation of the heater strip.

### Separation efficiency of different filter trenches

Three characteristics of the filter trench can affect the separation efficiency, namely the depth, geometry, and its position in relation to the channel. We examined the performance by changing one factor at a time to determine the optimal parameters for our system. Although 10 μL of the sample was sufficient for our device, to complete the experiments, we used 15 μL of whole blood for examination. The geometry of the filter trench is cylindrical, and the diameter was fixed at 2 mm.

#### (A) Effect of depth of filter trench

The depths of filter trench tested were 2 mm, 2.5 mm, and 3 mm, which could be obtained by varying the thickness of the PDMS slab. Because the relationship between the flow rate and depth could also affect separation efficiency, we examined the depth factor with the different flow rates, namely 2 μl min^−1^, 1 μl min^−1^, and 0.5 μL min^−1^. The result shown in Fig. 3 indicates that in our device, the separation efficiency is directly proportional to the depth of the trench and inversely proportional to the flow rate. Gravity exerted a significant effect on blood cells in the filter trench; therefore, an increase in the depth enhanced the sedimentation effect. In our study, the separation time was the duration from the time of entry of the sample into the trench to the time that the filtered plasma reached the biomarker test zone. The highest separation efficiency was observed at 0.5 μL min^−1^; however, the separation time was >10 min. Therefore, we considered 1 μL min^−1^, which had separation time of 4 min, the optimum value for flow rate for the device to meet the expectation of rapid operation. Therefore, our subsequent work involved improving the separation efficiency of the device by considering other aspects. In summary, the optimum depth of the filter trench and flow rate in this study were 3 mm and 1 μL min^−1^, respectively.

**Fig. 3.**
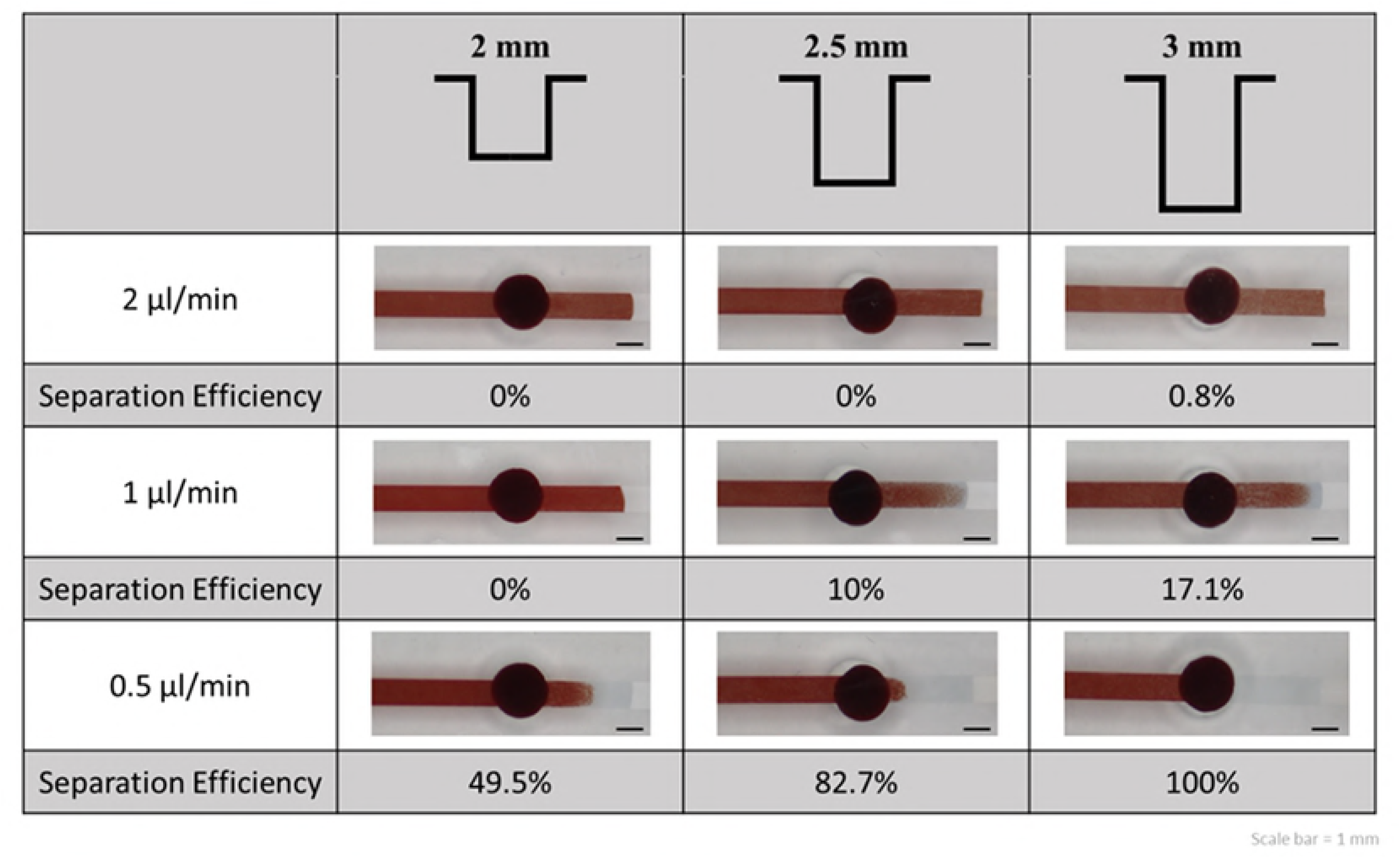
Separation efficiency at different depths of filter trench and flow rates

#### (B) Effect of geometry of filter trench

At different tilt angles (−45°, 0°, +45°), the cylindrical trench exhibited widely varying separation efficiencies (Fig. 4). A positive angle indicated that the cylinder was tilted toward the outlet, whereas a negative angle indicated that it was tilted toward the inlet. The effect of tilt angle on separation efficiency can be explained through flow simulation. When a blood cell or particle is suspended in the filter trench, it is affected by the buoyancy-corrected gravitational sedimentation force (*F*_*gb*_) and the fluid drag force (*F*_*d*_) (ESI† Fig. S1). The vertices of the curve indicate that *F*_*d*_ is 0; therefore, the particle is stationary in the fluid, and only *F*_*gb*_ acts on it. This is a critical stage for plasma separation because the effect of gravity is significant on blood cells in a low-speed area. Therefore, a forward tilted (+45°) structure showed higher efficiency than the original structure because the blood cells reached the low-speed area earlier, and the separation was completed effectively. In the backward tilted structure, turbulent flows interfered with the sedimentation of blood cells; therefore, the separation of plasma was difficult. We further tested the performance in three different geometries of filter trench, namely cylindrical, triangular prismatic, and trapezoidal columnar (ESI† Fig. S2). However, because of the difficulty of fabrication, the cylindrical geometry was the first option for trench design.

**Fig. 4.**
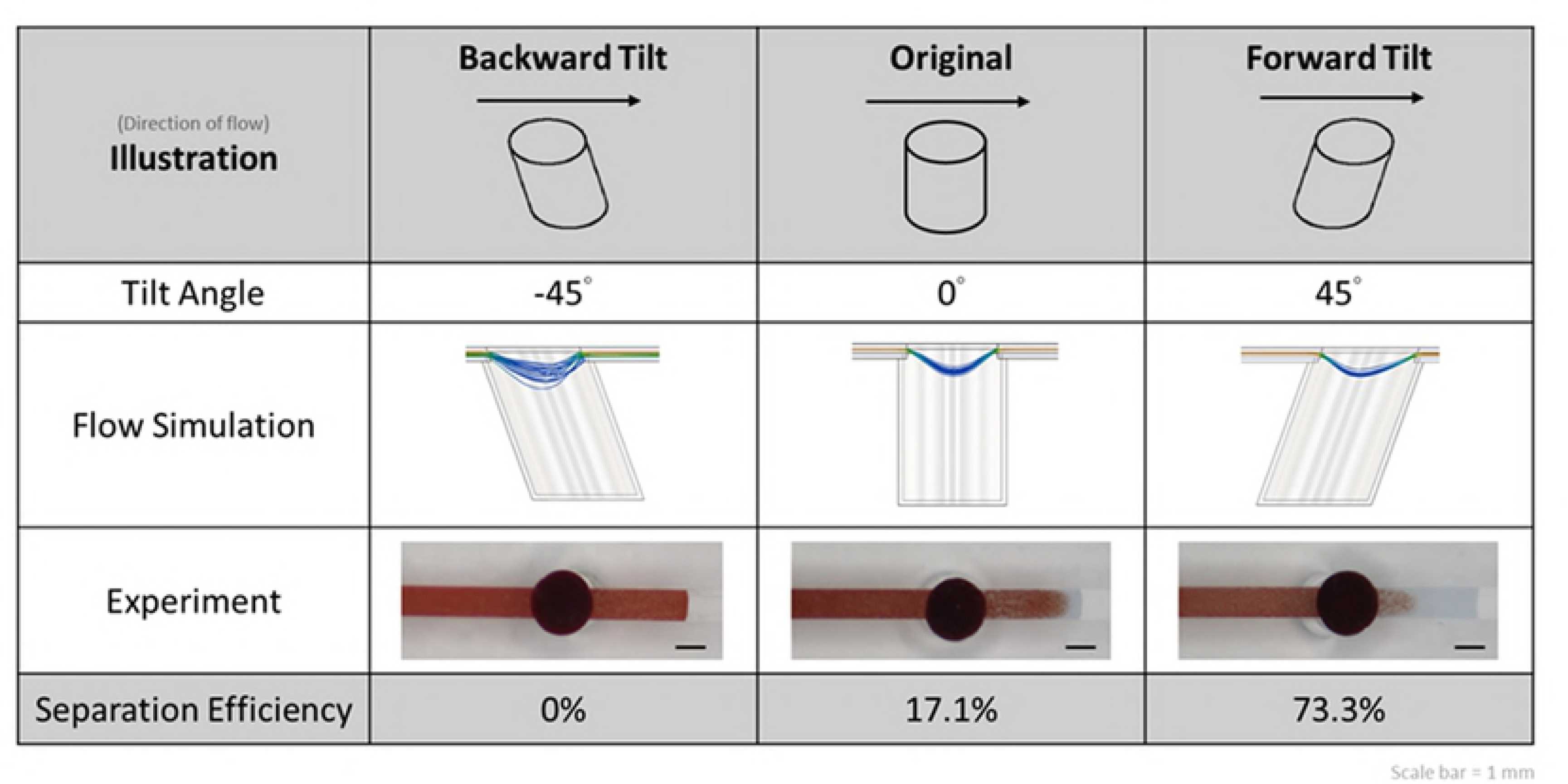
Separation efficiency with different geometries and results of flow simulation.

#### (C) Effect of relative positions of filter trench and channel

We examined the effect of the relative positions of the filter trench and channel on separation efficiencies. The channel was identified as buried or suspended if it was above or inside the trench (in a cross-sectional view), respectively. The forward-tilted trench showed a considerably higher separation efficiency than the original trench (73.3% and 17.1%, respectively) (Fig. 5A). However, a combination of the forward-tilted trench and suspended channel further increased the separation efficiency to 99.7%. Although the hematocrit is a crucial factor affecting the separation efficiency, the forward-tilted trench with suspended channel consistently exhibited the highest separation efficiency in each repeat experiment (data not shown).

**Fig. 5.**
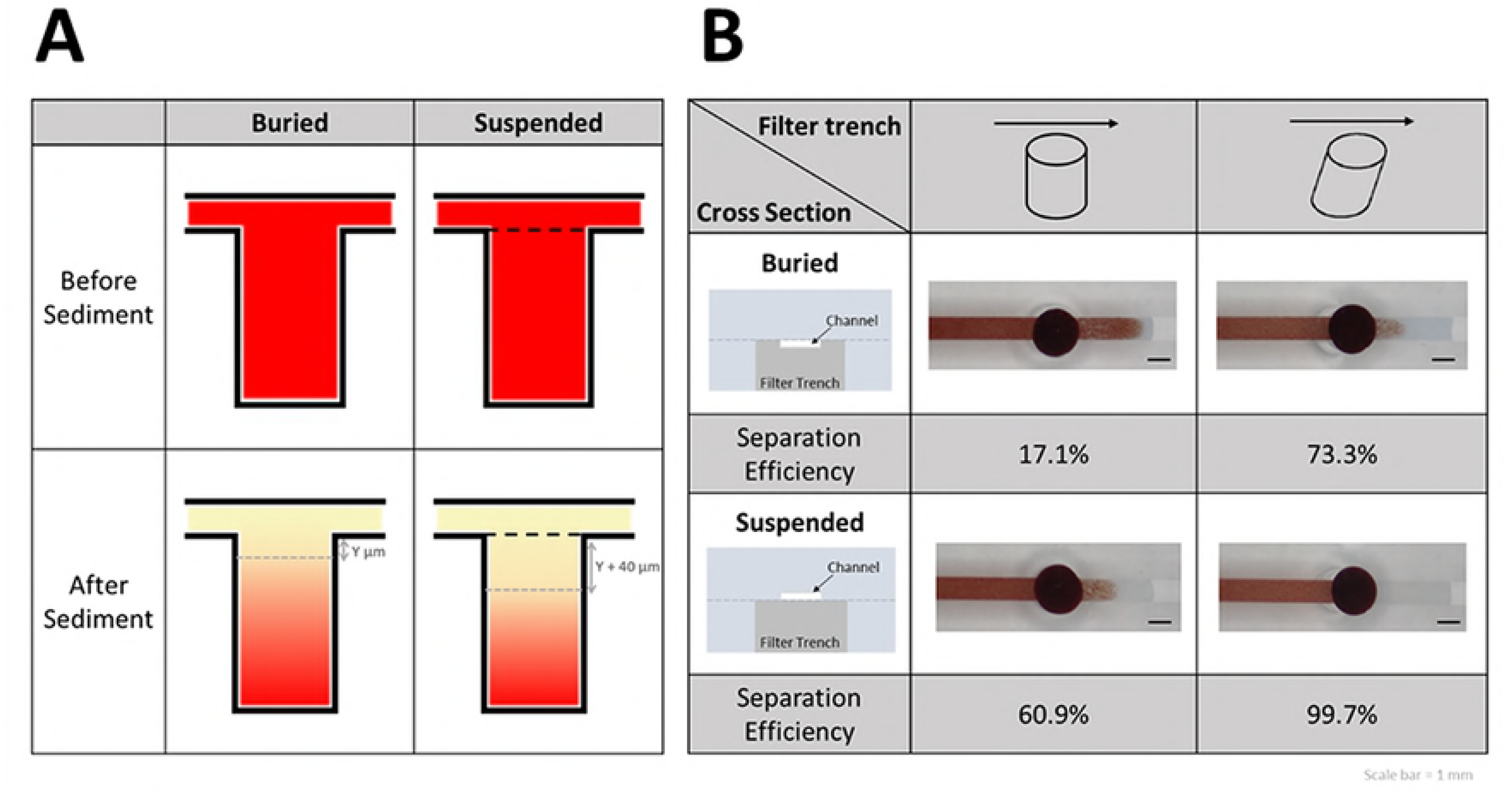
(A) Separation efficiencies obtained using the buried and suspended channel designs analyzed using flow simulation. (B) Relative positions of the channel and filter trench affect separation efficiency.

Suppose that the whole-blood samples that flowed into the buried or suspended channel were identical in volume and hematocrit, and thus, the amounts of sedimented RBCs were identical. In the suspended channel design, the channel was located above the filter trench; consequently, the distance between the surface of sedimented RBCs and the microchannel was greater than that in the buried design. The relatively great distance protected the sedimented RBCs from the interference of turbulent flows (Fig. 5B). This physical phenomenon caused the RBCs to settle at the bottom of the trench without being driven upward by the turbulent flows. Consequently, a higher separation efficiency was obtained with the suspended channel, and high-purity plasma could be collected at the outlet of the filter trench.

### Thermopneumatic microfluidic suction system

A microfluidic chip is highly suited for achieving POCT because it integrates multiple tedious laboratory techniques into a tiny chip. However, a proper force mechanism has been the major challenge to actual application. Recently, several research groups have adopted the principle of differential pressure for operating microfluidic devices without external pumps; however, portability and commercialization have not been achieved because of the difficulties involved in creating and maintaining low-pressure environments.

The use of the temperature of air to affect air density is a simple but effective method for creating and maintaining low-pressure environments. We applied the heater strip with the microchannel to achieve the same effect as a vacuum generator. Moreover, the power source of the heater strip was a pair of dry cells, which are inexpensive, portable, and reusable. Thus, a low-pressure environment can be easily created anytime and anywhere. The flow rate (*Q*_*flow*_) is defined as the ratio of volume change (*ΔV*) over time (*Δt*). If the temperature change is related to a constant cooling rate *k* by *ΔT = k*^⋅^*Δt*, the flow rate can be obtained as follows:

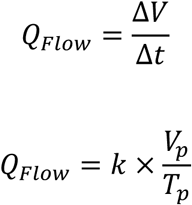

where *k* is constant, *V*_*p*_ is the volume of pressure chamber, and *T*_*p*_ is the temperature of pressure chamber. Therefore, the flow has almost no pulsation during the stroke, and precise temperature control is achieved.

Our device had an inlet and a straight channel with a filter trench in the center, and the end of channel was connected to a suction chamber (Fig. 1). The heater strip was folded into a square-wave-like structure and installed immediately under the chamber to heat the air in the chamber uniformly. The visible and thermal images of the heating process and elapsed time are shown in Fig. 6A. Because of the structural properties of the microenvironment, a slight change in temperature was effective for sample suction. An increase of only 4°C was sufficient for device operation (Fig. 6). Most importantly, this heating mechanism only affected the chamber zone during the entire process; therefore, the risk of heat denaturation of the sample, biomarkers, or analytes was negligible. In addition, flow control is a critical factor affecting separation efficiency. Therefore, optimizing the flow rate by adjusting the size of chamber, the length of heater strip, and the heating time is essential for this force mechanism. Many pumping applications, such as filling a biochemical microreactor chip, do not require constant flow rates, and ensuring that a defined volume of liquid is transported into the measurement cell within a given time is sufficient. However, for blood separation by using a gravity sedimentation system, the ideal flow rate is either constant or slightly quick in the beginning and gently slowing down as the blood sample reaches the filter trench zone. A stable flow rate is essential to prevent flow disturbance in the filter trench wells, which may affect the sedimentation of blood cells. This requirement is highly suited to the characteristics of a thermopneumatic microfluidic suction system.

**Fig. 6.**
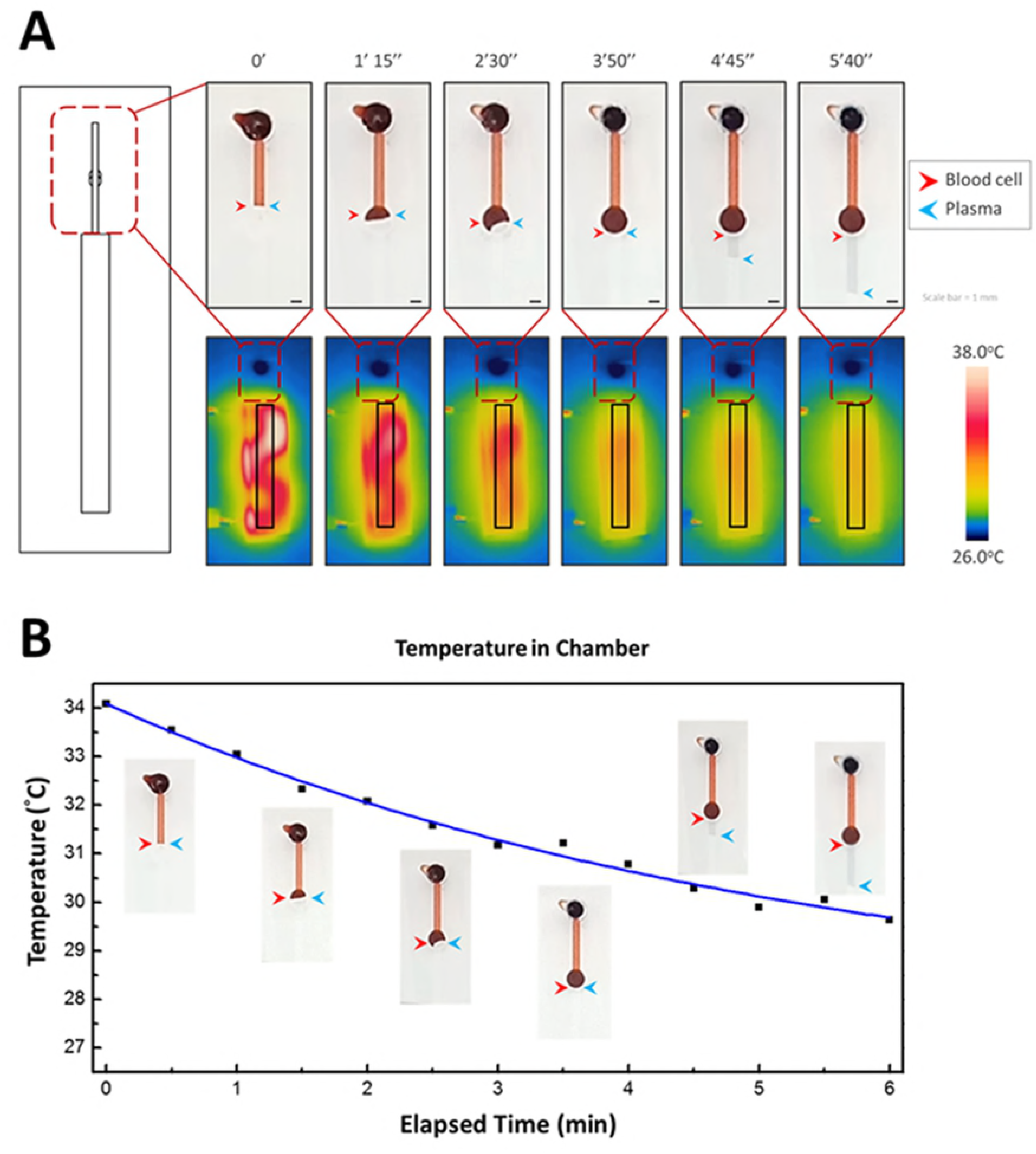
(A) Visible and thermal images of the microfluidic device during heating. The heater strip was folded into square-wave-like shape and installed immediately under the suction chamber to effectively heat the air inside the chamber. The red and blue arrows indicate the blood cells and plasma, respectively. The chromaticity bar indicates the temperature from 26°C (room temperature) to 38°C. The system works effectively with relatively small variations in temperature. (B) Chart of temperature reduction inside the chamber, and six actual images inserted near the curves indicate the stages of separation. The chart starts from the cooling down (i.e., from 2 min). The whole-blood sample was sucked into the channel immediately after loading. At 30 s after loading, the sample reached the inlet of the filter trench and filled the entire trench at 6 min. At 8 min, the filtered plasma entered the detection zone, and biomarker examination could be executed immediately.

### Thermopneumatic microfluidic suction system

Considering that POCT devices focus on cost-effective and rapid diagnosis rather than precise analysis, we demonstrated the concept of biomarker detection by using urine test strips (protein, glucose, and pH). The first step was to examine the color reaction. When a whole-blood sample was used, the strips exhibited a reddish-brown color because of the interference of RBCs; consequently detecting a color change was difficult. However, when a filtered plasma sample that was transparent was used, color reactions were easily detected (protein, glucose, and pH) (ESI† Fig. S4). We designed a notch near the outlet of filter trench and installed a urine test paper inside it. The color reaction was sufficiently clear for observation, which indicated that the plasma was effectively separated by our system and the biomarker detection test was completed (Fig. 7). The chart of temperature variation in the detection zone of the microfluidic device shows that the temperature change was <0.5°C when the isolated plasma was sucked into the detection zone (blue area). Therefore, the sample or analytes were denatured by the heating-force mechanism (ESI† Fig. S5). This device has potential to realize POCT in the near future because of its low cost, easy fabrication, simple two-step operation, and portability.

**Fig. 7.**
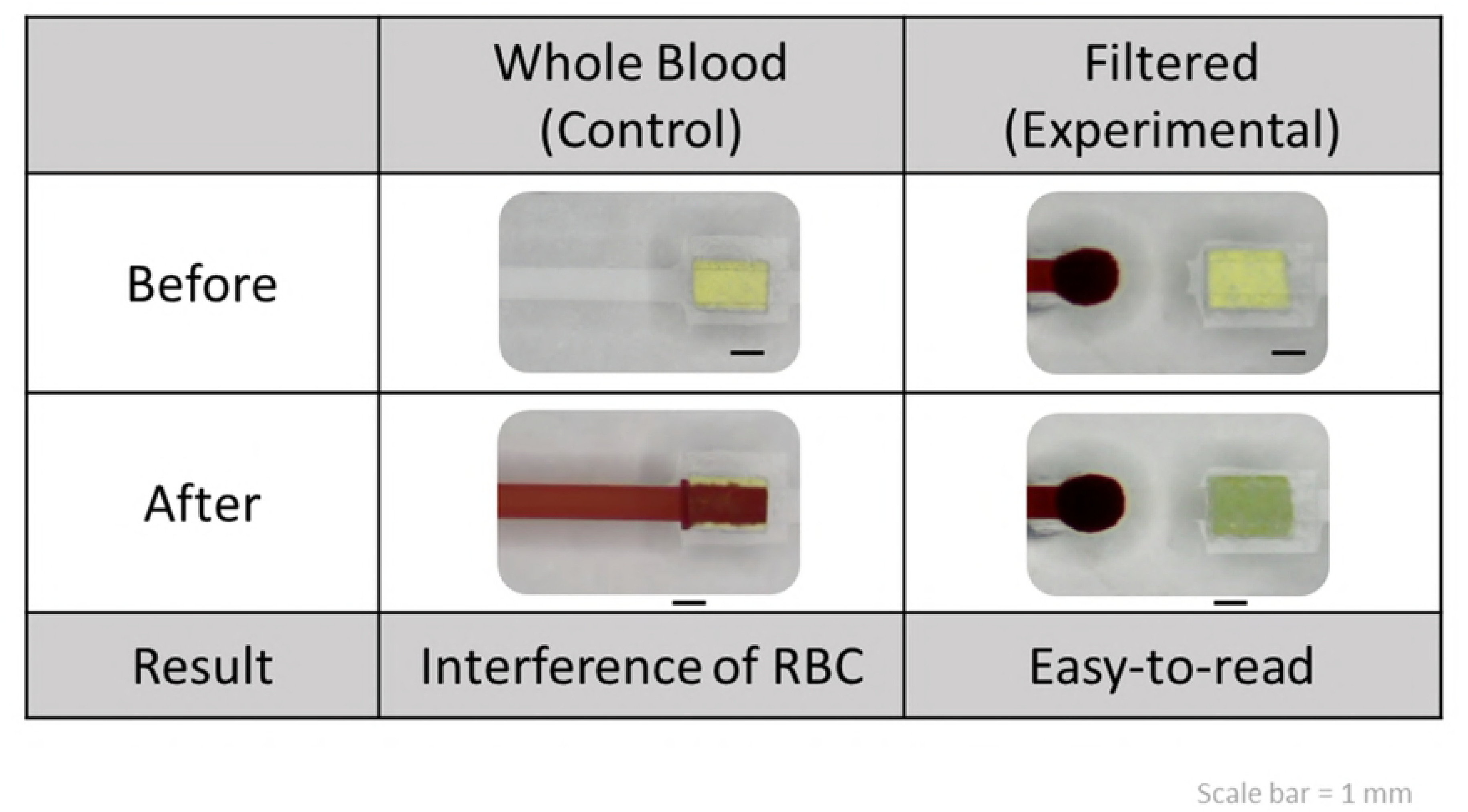
Comparison between the whole blood and filtered plasma samples on devices with and without a filter trench in the middle of the channel. The device without a filter trench used whole blood for detection and showed a result with RBC interference. However, the device with a filter trench separated the transparent plasma; thus, the color change can be observed easily and clearly.

## Conclusion

A portable, rapid, easy-to-fabricate, and easy-to-use microfluidic chip was designed for separating plasma from whole-blood samples. In this study, we examined the separation efficiency from many aspects, such as the design of the microchannel, filter trench geometry, and relative positions of the filter trench and channel. After parameter optimization, the separation efficiency was enhanced from 17.1% to 99.7%. In addition, the blood separation device and self-strained system were integrated in the chip. The separation required a heating mechanism that created a low-pressure environment and separated the plasma within 2 min. The proposed device reduced the entire working time to 6 min and yielded >1 μL of isolated plasma, which is sufficient for most bioassays (Table 1). Finally, the cost of consumables with the device was as low as 1 USD, which demonstrated its considerable potential to realize the concept of POCT in the near future.

**Table 1.** Blood cell separation microdevices and their performance

| Method | Blood sample ($\mu\text{l}$) | Isolated plasma ($\mu\text{l}$) | Separation Time (min) | Sample driving force | Ref. |
| --- | --- | --- | --- | --- | --- |
| Sedimentation on chip | 5 | 0.4 | 10 | Vacuum suction | [27] |
| Filtration membrane | 3 | 0.5 | 20 | Vacuum suction | [28] |
| Micropost array | 5 | 0.2 | 5 | Capillary force | [29] |
| Micropillar array | 20 | 0.2 | 3 | Capillary force | [30] |
| hydrophobic patch | 3 | 0.2 | 10 | Capillary force | [31] |
| hydrophobic patch | 4 | 0.4 | 15 | Capillary force | [32] |
| Sedimentation on chip | 10 | 0.8 | 15 | Capillary force | [33] |
| Sedimentation on chip | 10 | 2 | 6 | Thermo pneumatic suction | This work |

## Acknowledgements

This study was financially supported by the Ministry of Science and Technology (MOST) of Taiwan (106-2113-M-009-013-MY2) and Center for Emergent Functional Matter Science of National Chiao Tung University, Featured Areas Research Center Program, within the framework of the Higher Education Sprout Project of the Ministry of Education (MOE) in Taiwan.

## Author contributions

Dr. BR Li was director of the project. Ms. CH Yang performed most experiments. Ms. YL Hsien organized and wrote the paper, Dr. PH Tsou discussed and helped to obtain human blood samples from healthy donors according to a protocol permitted by the Institutional Review Board (IRB).

## Supporting Information

**Fig. S1** Modeling the principle force acting on a suspended particle.

**Fig. S2** Sparation efficiency in different geometries.

**Fig. S3** Illustration of the cross section of the buried and suspended channels.

**Fig. S4** Bioassays with RBCs contained and removed sample on test paper.

**Fig. S5** Temperature variation in the detection zone of the microfluidic chip.

